# Development of a Spectral Flow Cytometry Analysis Pipeline for High-Dimensional Immune Cell Characterization

**DOI:** 10.1101/2024.06.19.599633

**Authors:** Donald Vardaman, Md Akkas Ali, Chase Bolding, Harrison Tidwell, Holly Stephens, Daniel J. Tyrrell

## Abstract

Flow cytometry is a widely used technique for immune cell analysis, offering insights into cell composition and function. Spectral flow cytometry allows for high-dimensional analysis of immune cells, overcoming limitations of conventional flow cytometry. However, analyzing data from large antibody panels can be challenging using traditional bi-axial gating strategies. Here, we present a novel analysis pipeline designed to improve analysis of spectral flow cytometry. We employ this method to identify rare T cell populations in aging. We isolated splenocytes from young (2–3 months) and aged (18–19 months) female mice then stained these with a panel of 20 fluorescently labeled antibodies. Spectral flow cytometry was performed, followed by data processing and analysis using Python within a Jupyter Notebook environment to perform batch correction, unsupervised clustering, dimensionality reduction, and differential expression analysis. Our analysis of 3,776,804 T cells from 11 spleens revealed 34 distinct T cell clusters identified by surface marker expression. We observed significant differences between young and aged mice, with certain clusters enriched in one age group over the other. Naïve, effector memory, and central memory CD8^+^ and CD4^+^ T cell subsets exhibited age-associated changes in abundance and marker expression. Additionally, γδ T cell clusters showed differential abundance between age groups. By leveraging high-dimensional analysis methods borrowed from single-cell RNA sequencing analysis, we identified age-related differences in T cell subsets, providing insights into the immune aging process. This approach offers a robust, free, and easily implemented analysis pipeline for spectral flow cytometry data that may facilitate the discovery of novel therapeutic targets for age-related immune dysfunction.

## Introduction

Flow cytometry is a widely and frequently used technique to explore immune cell composition and function in many fields of science and in clinical practice.^1–3^ Flow cytometry is a powerful approach to determine cell size, granularity, number, proportion, and expression of surface and intracellular proteins.^4–6^ In recent years, advances in flow cytometers and spectral computation has enabled spectral flow cytometry for high-dimensional analysis of immune cells.^7,8^ Spectral flow cytometry offers advantages in resolution, multiplexing, and quantitative accuracy. A major advantage over conventional flow cytometry is the ability to use more antibodies within the same staining panel to characterize distinct and rare populations of cells.^9^

Samples are typically split and stained with distinct panels of antibodies to characterize certain subsets of cells such as myeloid cells or lymphocytes. Splitting samples reduces the number of cells that can be profiled per panel, which can limit overall data production, especially where tissue is limited. Even by using a targeted panel of antibodies for specific cell types, inclusion of certain antibodies may be limited due to spectral overlap which occurs more easily in conventional flow cytometry.^8,10^

In conventional flow cytometry, fluorochromes are detected using a series of optical filters and detectors, each tuned to detect specific emission wavelengths.^11–14^ The filters have limitations on spectral resolution and the number of distinct fluorochromes that can be resolved simultaneously. In contrast, spectral flow cytometry captures the full emission spectra of fluorochromes.^7,8^ This is coupled with spectral unmixing algorithms to deconvolute the mixed spectra and assign each component to its respective fluorochrome.^15,16^ This allows for improved resolution and discrimination of closely overlapping signals, and thus more fluorochrome-linked antibodies can be used.

In conventional flow cytometry, antibody panels incorporating a live/dead stain typically include between 6-12 antibodies and are now able to scale up to 28 antibodies with optimization.^17^ As an example, a typical strategy for analyzing CD8^+^ T cells would involve gating on live, CD45^+^ CD3e^+^ CD8a^+^ cells.^17^ After this, surface markers such as CD62L, CD44, PD1, Tox, CD27, and CD28 may be used to further categorize distinct subsets of CD8^+^ T cells.^17,18^ Conventional flow cytometry gating strategies examine two markers at a time to subset cells.^19^ These subsets can then be drilled down to examine two more markers until all markers have been analyzed. If a flow cytometry panel has only six antibodies, there are only six possible unique gating sequences possible. All combinations are not typically considered because prior knowledge informs the gating strategy order.^17,20–23^

Spectral flow cytometry allows for additional antibodies to be used within the same panel even up to 50 colors in a single panel.^9,24,24^ By expanding the number of additional markers (i.e., CD49d, CXCR3, CCR5, CXCR6, CD44, CD27, TCRY, CD28, PD1, LAG3, KLRG1, CD62L, MHCII, MHCI, TOX, and B220), the complexity for gating strategies increases substantially. While it is possible to rely on hypothesis-based or knowledge-driven gating strategies, the addition of several antibodies increases the number of gating steps, thus making analyses notably more difficult to determine which markers to use in combination and which order to examine them in. Imprecise or sub-optimal gating strategies could impact the number of cell populations found in a sample. Spectral flow cytometry enables the simultaneous detection of a larger number of markers in a single panel and a more comprehensive understanding of cell populations; however, data analysis can be cumbersome with conventional gating strategies.

Here, we introduce a novel spectral flow cytometry panel and data analysis pipeline that incorporates aspects of conventional flow cytometry gating with unsupervised clustering and analysis borrowed from single-cell RNA sequencing analysis. We demonstrate the feasibility and utility of this approach with minimal coding input by using Jupyter notebooks. We used this pipeline to identify differences in CD8^+^ and CD4^+^ T cell subsets between young and old splenic T cells. This procedure allows for high dimensional analysis of abundant and rare cell populations.

## Methods

### Data availability

Raw data including flow cytometry .fcs files and .wsp file along with flow cytometry scale values used to run the pipeline (.csv file), outputs from the pipeline, and data used to generate figures are available on Figshare at https://figshare.com/projects/Development_of_a_Spectral_Flow_Cytometry_Analysis_Pipeline_for_High-Dimensional_Immune_Cell_Characterization/210475. Code is available on Github at https://github.com/mdakkasali/Spectral_Flow_Cytometry_Data_Analysis. Data and code used for Supplemental Figures are available from the authors upon reasonable request.

### Study approval

All animal experiments were carried out in accordance with the Institutional Animal Care and Use Committee at the University of Alabama at Birmingham (protocol no. 22627).

### Mice and diet

C57BL/6N WT young (2–3_months) and aged (18–19_months) female mice were obtained from the National Institute on Aging rodent colony and Charles River Breeding Laboratories (stock no. 027). All mice were maintained on a 12-h light/dark cycle with free access to food and water. Sizes of experimental groups were based on power calculation from our previous studies.

### Cell isolation

Spleens were harvested postmortem and transferred to a cell strainer (70µm) on a 50mL tube. Each spleen was cut into 4 equal sections on the strainer. Using the rubber end of a sterile syringe plunger, the spleen was smushed through the strainer mesh. The strainer was rinsed with 30mL of 1x D-PBS. The sample was then centrifuged at 500xG for 10 minutes at 25 degrees C. The supernatant was discarded and the pellet was resuspended in 4mL of 1x D-PBS and vortexed briefly at medium speed. Cells were resuspended in 500µL and passed through a cap strainer mesh (70µm) into a 5mL round-bottom FACS tube. The samples were centrifuged again at 500xG for 5 minutes at 25 degrees C. The supernatant was discarded, the pellet was vortexed briefly at medium speed, and staining protocol commenced.

For cell isolation from the brain, we perfused mice with PBS and isolated the brain. The brain was minced and incubated for 20 minutes in 2mL of enzyme mixture (1g collagenase IV and 20 mg DNAse I in 500 mL of media was prepared) at 37°C. The digested tissue was strained through a 70µm cell strainer into a clean 50mL conical tube, rinsed with cold PBS (Ca²⁺ and Mg²⁺-free) up to 40mL, and centrifuged at 500xg for 8 minutes at 4°C. The supernatant was discarded, and the pellet was resuspended in 7 mL of media and mixed with 3 ml of 90% Percoll, then gently vortexed. 2-3mL of 70% percoll solution was injected beneath the 90% Percoll solution. The brain-media mixture was centrifuged for 32 minutes and the buffy coat was extracted. The isolated cells were washed with cold PBS, centrifuged, resuspended in 500µL of cold PBS, transferred through 70µm cell strainer into a 5mL FACS tube for subsequent antibody staining.

### Spectral flow cytometry

Single-cell suspensions in residual D-PBS in 5mL FACS tubes were incubated with 20µL of Aqua Live/Dead stain diluted 1:100 in 1x D-PBS for 5 minutes at 25 degrees C. 50µL of anti-mouse CD16/32 Fc receptor blocking solution (1:50 in FACS buffer) was added to each sample and incubated for 10 minutes at 25 degrees C. Then 30µL of antibody master mix was added to each sample and incubated for 30 minutes at 25 degrees C. After incubation, each sample was washed with 2mL of FACS buffer and centrifuged at 500xG for 5 minutes 25 degrees C. The supernatant was discarded and the cell pellet was broken up by vortexing briefly at medium speed. Samples were incubated with 1mL of 4% paraformaldehyde for 15 minutes at 25 degrees C. Samples were washed two times and then resuspended in 200µL of FACS buffer and stored at 4 degrees C until running on the flow cytometer. Spectral flow cytometry was performed on BD FACSymphony A5 SE Cell Analyzer with BD FACSDiva Software version 9.0. Initial data processing was done with FlowJo version 10.10.

### Analysis pipeline

For computational analyses, we employed a Jupyter Notebook environment (version 1.5.1). This environment was configured on a virtual server infrastructure that allocated resources as per the requirements of our computational tasks. We ensured an appropriate setup for our analyses by loading specific environment modules necessary for our software dependencies. We utilized 32 CPUs to allow for parallel processing of our data-intensive tasks, optimizing computational time. Each CPU was allocated 10 GB of memory, totaling 320 GB of RAM, to ensure sufficient memory for our processes and to prevent bottlenecks during data analysis. Our computational jobs were set to run for a maximum of 30 hours to ensure the completion of long-running processes without interruption. We selected the medium partition for our job submission, balancing the need for resources with availability within our institutional computational cluster.

Analysis was performed using Python (version 3.12.0) within a Jupyter Notebook (version 1.5.1) framework to ensure reproducibility. We used Python to analyze single-cell data from spectral flow cytometry. Specifically, we focused on surface protein expression used for staining cells. All flow cytometry data were initially analyzed using FlowJo (Version 10.9.0). We first gated out the debris, then gated on singlets, then live cells, then CD45+ cells, then CD3e+ and B220-cells to isolate all T cells. From this population of T cells, we exported the raw channel values and scaled values as CSV files.

We loaded these data into a Jupyter Notebook as a single dataframe using the ‘glob’ and ‘pandas’ packages for further analysis. The main dataset was subsequently converted into an AnnData object with corresponding metadata using the ‘sc.AnnData(expr_data)’ function with metadata such as sample identification and group identification using ‘adata.obs = metadata’. Technical noise and batch effects were addressed using the Scanpy Z-score scaling of cell-level data using the ‘sc.pp.scale()’ function followed by ComBat algorithm ‘sc.pp.combat()’ on the ‘Group’ variable for batch correction.^25,26^

We then performed dimensionality reduction using principal component analysis using ‘sc.tl.pca()’ function with ‘arpack’, enabling the subsequent construction of neighborhood graphs with ‘sc.pp.neighbors()’ using the PCA. Cells were plotted with uniform manifold approximation projection (UMAP) with ‘sc.tl.umap()’ then clusters were identified by Leiden algorithm using ‘sc.tl.leiden()’^27^, effectively identifying distinct cellular subpopulations.

For comparison of leiden to Phenograph, X-shift, and FlowSOM, we used the “phenograph”, “sklearn.cluster: DBSCAN”, and “minisom” python packages, respectively. The clustering was set to identify 20 clusters and these clusters were shown in UMAP. Silhouette score and Davies-Bouldin index were calculated for each of these using the “sklearn.metrics” python package. We also performed Leiden clustering on data that were not subjected to principle components analysis by performing “sc.pp.neighbors()” with “use_rep=’None’”.

Differential expression analysis was conducted to identify marker genes across distinct cell populations defined by unsupervised clustering (Leiden algorithm). We utilized the ‘rank_genes_groups’ function in Scanpy to conduct a t-test, identifying genes that were differentially expressed with statistical significance among the clusters defined by the ‘leiden’ label. After identifying marker genes, a dendrogram was computed to show hierarchical relationships between clusters based on expression profile. Data were represented via dot plot to represent expression level and percent of cells expressing top-ranked genes from each cluster. Genes were selected based on ranking in differential expression analysis. Plots were generated using the ‘dotplot’ function in Scanpy and ordered by dendrogram.

To ensure ease of use and reproducibility, our analysis process from raw data importation through to the generation of visual outputs is documented within a Jupyter Notebook and deposited on Github (https://github.com/mdakkasali/Spectral_Flow_Cytometry_Data_Analysis). This repository contains a tutorial for setting up the Python and Jupyter environment with appropriate packages to complete the analysis. The Jupyter Notebook itself contains detailed annotation to direct users to specific lines of code that need to be updated for user implementation (i.e. the folder path to where the user raw data files are located and leiden resolution appropriate for the type of data and number of cells). The raw data used in the current manuscript is also deposited on Github to enable users to test our analysis pipeline without updating or changing any code in the Jupyter Notebook.

## Results

An overview of our workflow and experimental design is shown in Figure 1. We stained splenocytes from 3-month and 18-month old mice with surface antibodies listed in Table 1 and performed spectral flow cytometry. We also performed spectral flow cytometry on single-stained bead samples. Individual spectral profiles for bead single-stained samples are shown in Supplemental Figure 1. In the 11 samples, we exported compensated “.fcs” files and performed initial gating on FlowJo software. We gated out debris, then gated on live singlet cells that expressed CD45 and CD3e (Supplemental Figure 2A). We then exported the scaled CD3e^+^ cell data from FlowJo as a concatenated “.csv” file and analyzed the expression of the remaining 16 fluorescent proteins using our analysis pipeline which we coded in Python.

**Figure 1:**
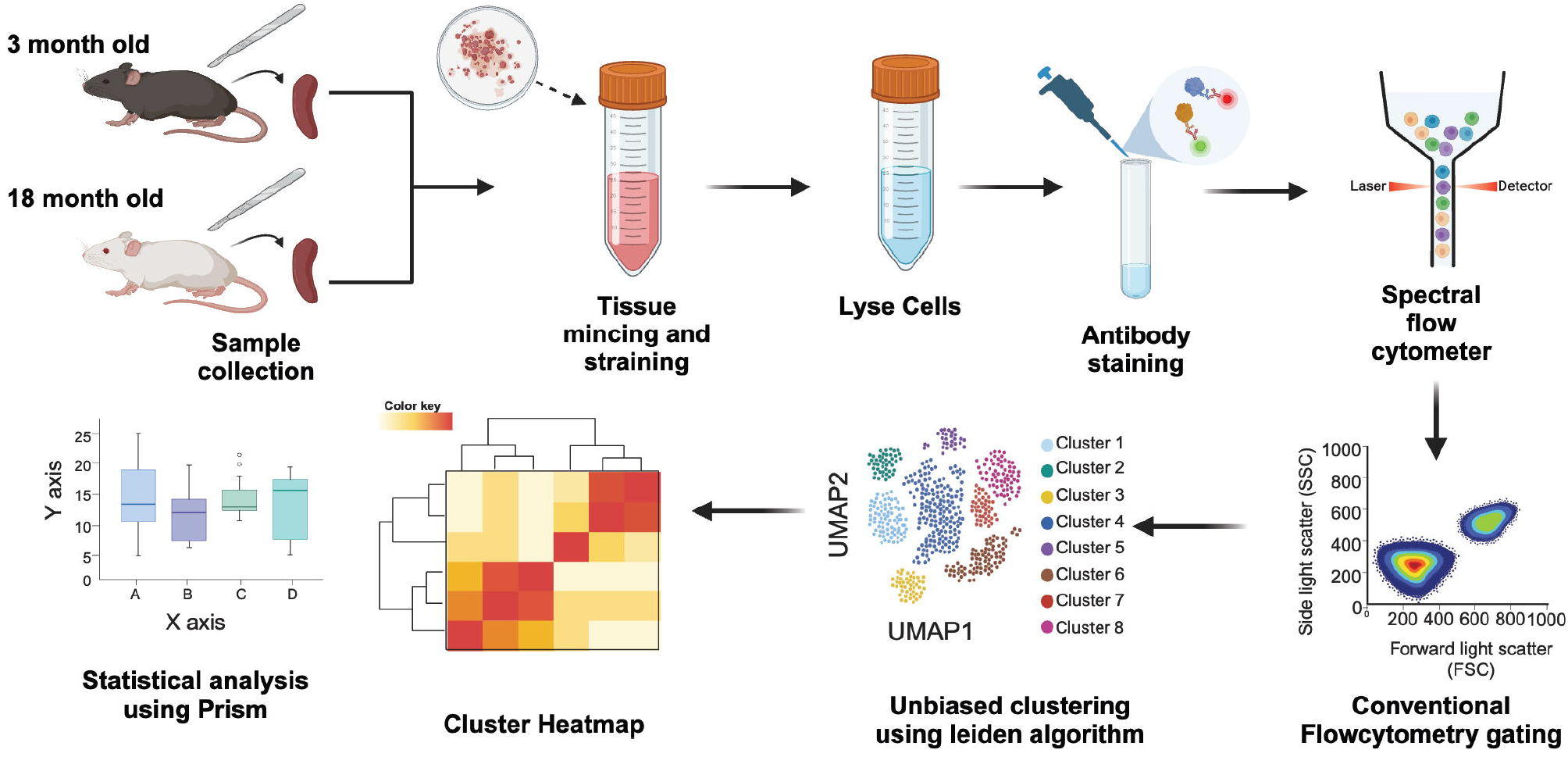
Overall experimental design. Diagram illustrating the workflow of spectral flow cytometry beginning with sample collection, tissue processing, cell staining, spectral flow cytometry, conventional gating on T cells, unbiased clustering and dimensional reduction, and data analysis.

**Table 1:**
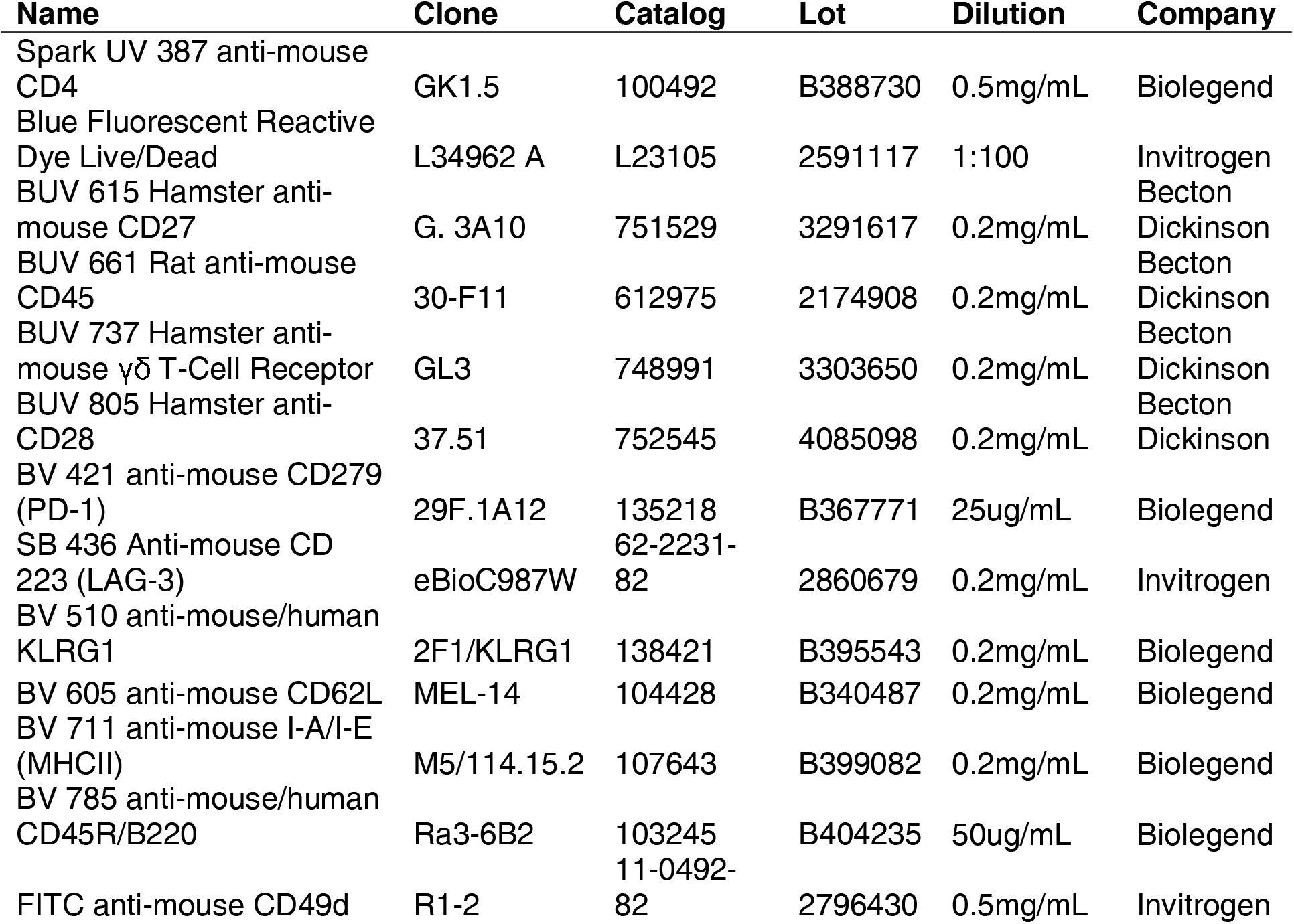
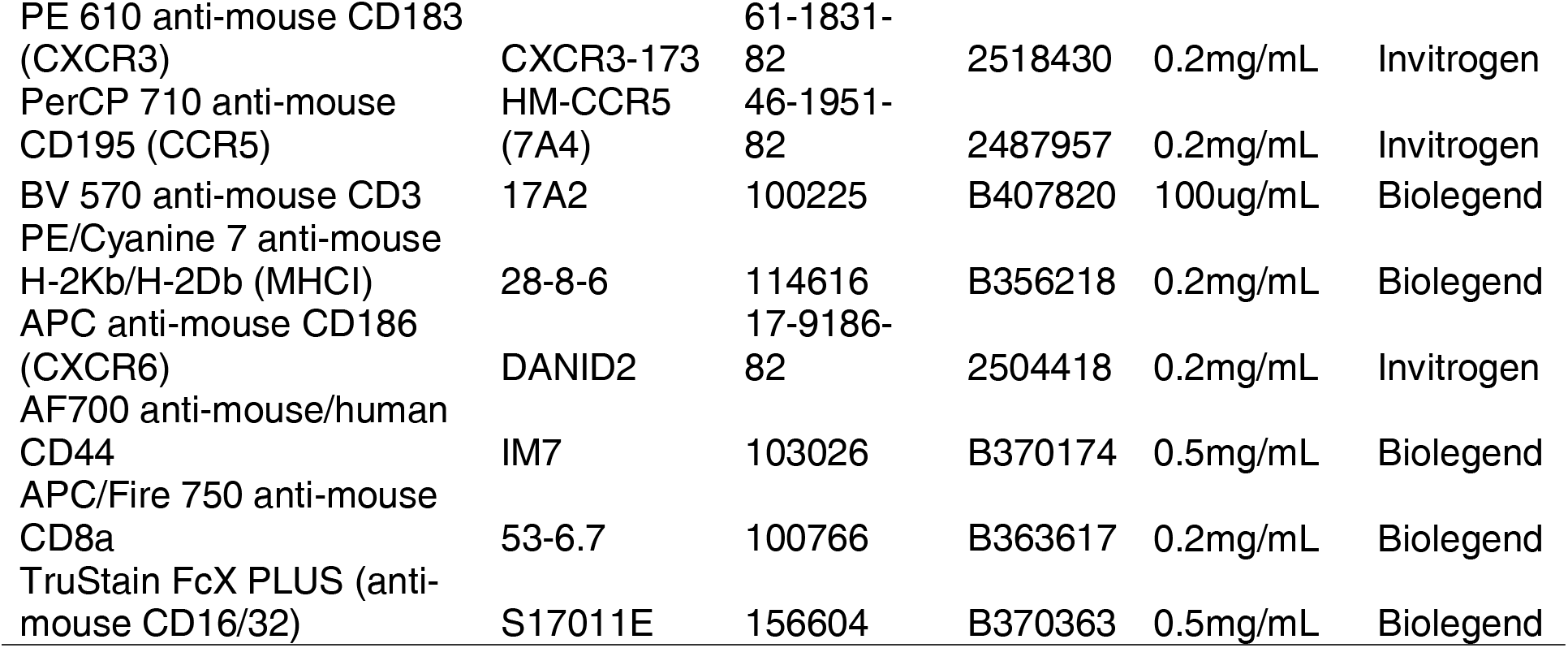
Antibodies used for flow cytometry staining.

We first performed batch correction, leiden algorithm^27^ for unsupervised clustering, and uniform manifold approximation projection (UMAP)^28^ for dimensionality reduction and plotting (Figure 1). Our analysis pipeline enables data analysis with the ability to customize antibody panels for various surface markers. We manually gated on live, CD45^+^ CD3e^+^ T cells and exported the scaled channel value data on the remaining antibody markers. This included 3,776,804 cells from 11 spleens (N=6x 3-month and N=5x 18-month) and 16 antibodies to phenotype T cells. After running our pipeline, we generated a UMAP to show the 34 distinct cell clusters identified by leiden algorithm (Figure 2A). This allows for a visual representation of the many different cell types identified with all cells from all animals visualized together. The leiden clusters ranged in size from the largest, Cluster 0, with 312,377 cells to the smallest, Cluster 34, with only 33 cells (Figure 1A). We also plotted the single-stained bead controls and unstained sample using UMAP to demonstrate how these samples cluster (Supplemental Figure 3A).

**Figure 2:**
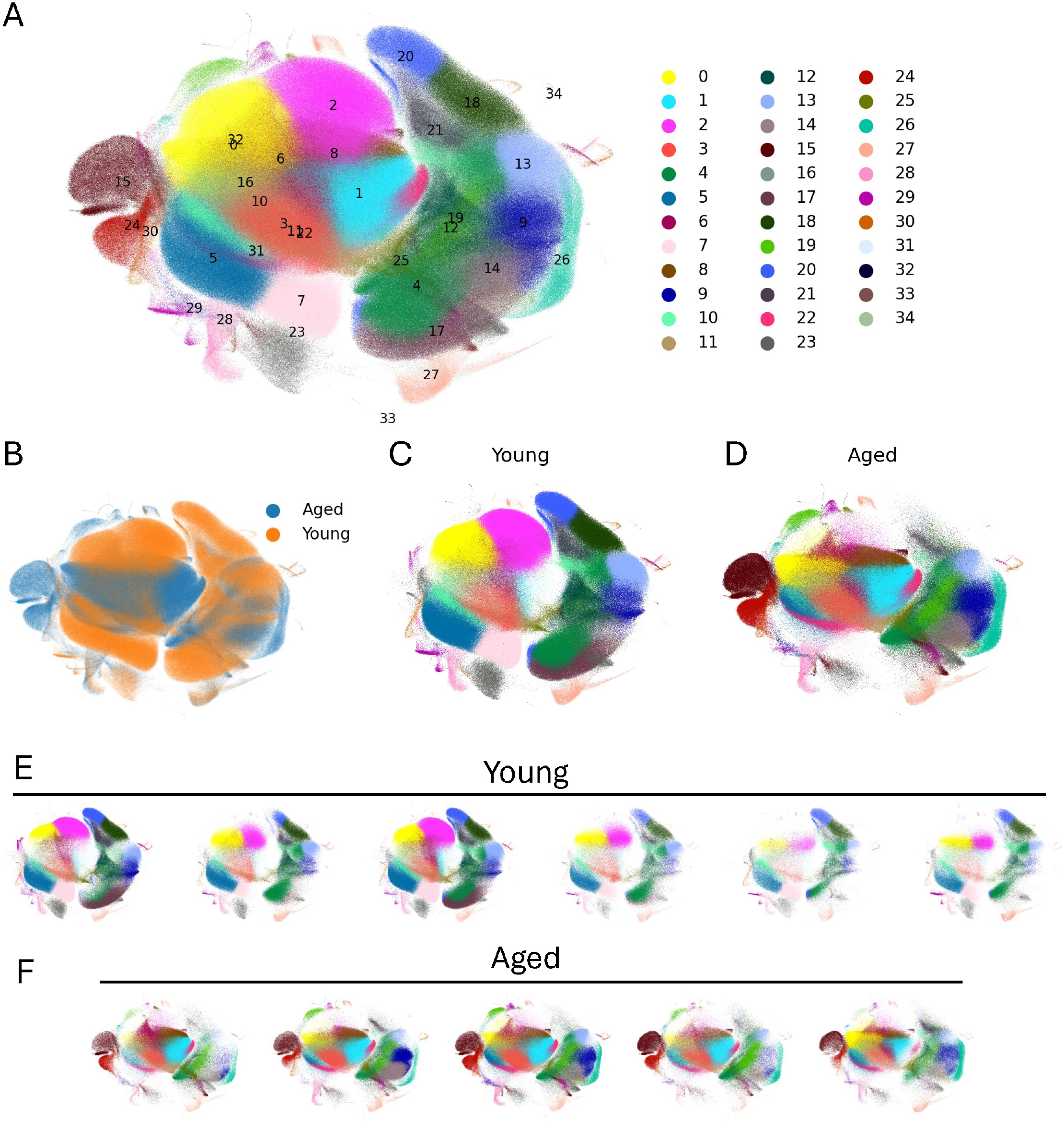
T cell clusters differ by age. A) Uniform manifold approximation projection (UMAP) plot showing the distribution of 3,776,804 cells categorized into 34 leiden clusters based on expression profiles of 16 immunological markers: CD49d, CXCR3, CCR5, CXCR6, CD44, CD8a, CD4, CD27, TCRY, CD28, PD1, LAG3, KLRG1, CD62L, MHCII, and MHCI. Each cluster is uniquely numbered and color-coded. B) UMAP showing the distribution of cells by age group. UMAPs showing the cells by leiden cluster in young (C) and aged (D) mice. UMAPs showing leiden clusters for all individual young (E) and aged (F) mice. N=6 mice at 3-months old and N=5 mice at 18-months of age. All mice are female C57BL/6N.

To compare the Leiden clustering algorithm with FlowSOM^29^, PhenoGraph^30^, and X-shift^31^, we used a smaller dataset of T cells and B cells isolated from the brain. We examined 69,555 cells from N=19 mice. We employed our analysis pipeline and identified 20 clusters by Leiden algorithm (Supplemental Figure 4A). We also performed the Leiden algorithm on the dataset without performing principle components analysis to preserve the high-dimensional structure of the data (Supplemental Figure 4B). This also resulted in 20 clusters. We then performed clustering with Phenograph, X-shift, and FlowSOM (Supplemental Figures 4C-E). Phenograph and FlowSOM resulted in 20 clusters of cells and X-shift resulted in 13 clusters. We then compared these methods using the Davies-Bouldin Index^32^ and Silhouette Score^33^. The Davies-Bouldin Index ranges from 0 to infinity with lower numbers representing greater partitioning. We found that the Leiden algorithm performed on PCA dimension-reduced spectral flow cytometry data provides the lowest Davies-Bouldin index (1.79) compared with X-shift (−3.210), Phenograph (1.91), and FlowSOM (2.10) (Supplemental Figure 4F). The Silhouette score ranges from −1 to 1 and is an index of how well individual cells (in this case) fit within their cluster where 1 is a perfect fit and −1 indicates an inappropriate cluster assignment. The Leiden clustering performed on PCA dimension-reduced data also resulted in the highest Silhouette score (0.093) compared with X-shift (1.88), Phenograph (0.089), and FlowSOM (0.075) (Supplemental Figure 4G). These data indicate that Leiden outperforms Phenograph, X-shift, and FlowSOM at appropriate clustering of spectral flow cytometry data.

Our splenic dataset included n=1,753,089 cells from young spleens and n=2,023,715 cells from aged. Some clusters were more represented in either young or aged samples (Figure 2B). Differences between age groups are apparent after stratifying the young and aged samples and coloring by leiden cluster (Figure 2B-C). Notably, clusters such as 2, 5, 7, 17, and 20 are present in the young samples but are nearly absent in the aged samples. In contrast, clusters 8, 15, 24, and 26 are nearly absent in young animals but are present in aged animals. Several clusters, such as 0 and 3 are represented in both young and aged groups. These differences are uniformly evident across each young (Figure 2E) and aged (Figure 2F) sample.

We next show the expression of individual surface proteins by dot-heatmap clustered by dendrogram (Figure 3A). We also plotted individual protein expression markers by UMAP, which demonstrates the clusters of cells expressing certain markers (Figure 3B). Based on these, we identified the cells on the right-hand side of the UMAP as CD8^+^ T cells and those on the left as mostly CD4^+^ T cells. Some clusters, like cluster 34, had a high expression of a single protein (i.e., CXCR3). While other clusters, such as Cluster 32, had a wide variety of protein expression. Using the dendrogram and UMAP together, we can identify cell types that make up each cluster. We also plotted our single-stained bead control samples by dot-heatmap (Supplemental Figure 3B) and by UMAP (Supplemental Figure 3C).

**Figure 3.**
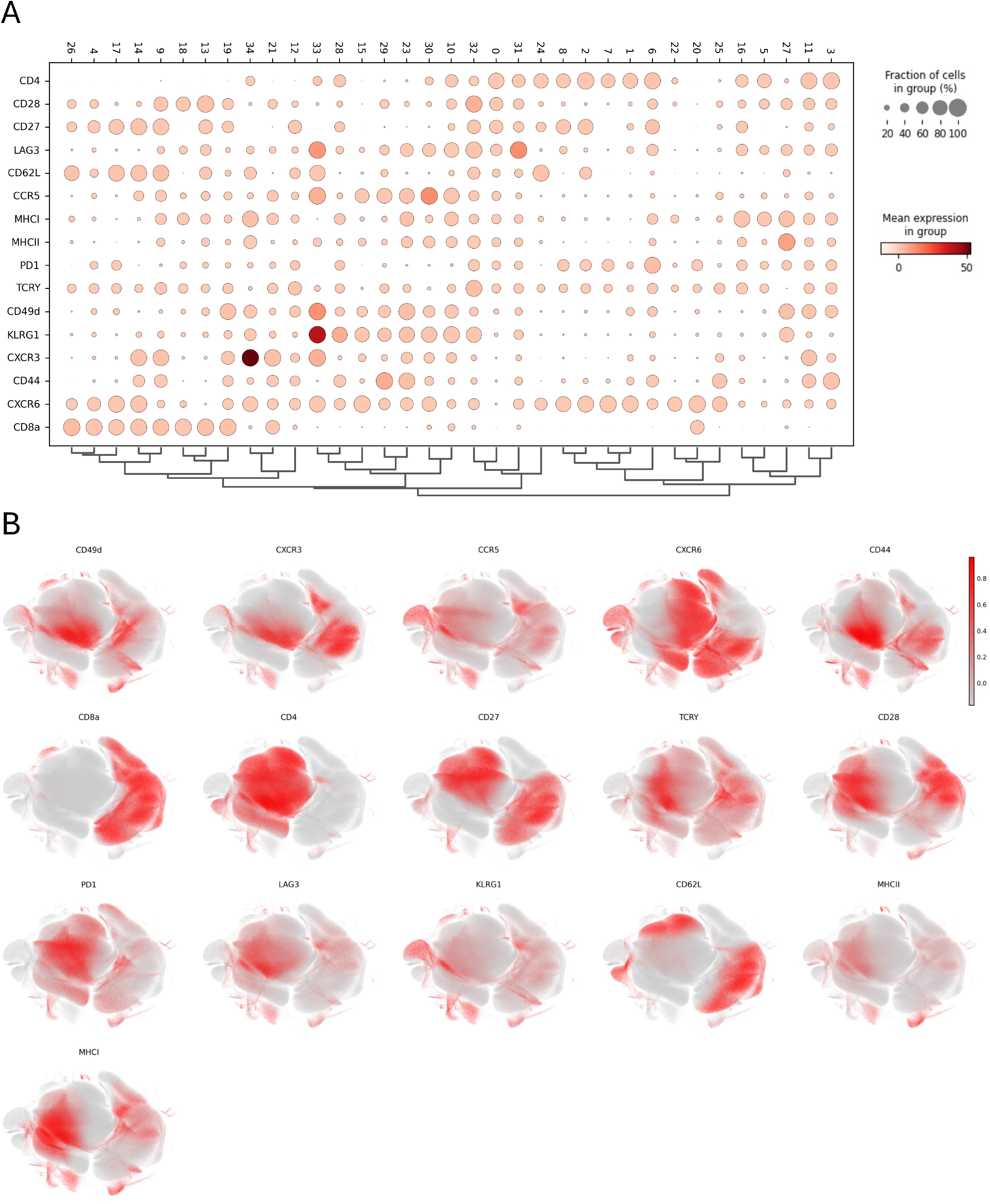
Marker protein expression by leiden cluster. A) Hierarchical clustering analysis reveals cellular subpopulations based on marker protein expression profiles and plotted as a dot-heatmap dendrogram. Expression is shown for ‘CD49d’, ‘CXCR3’, ‘CCR5’, ‘CXCR6’, ‘CD44’, ‘CD8a’, ‘CD4’, ‘CD27’, ‘TCRY’, ‘CD28’, ‘PD1’, ‘LAG3’, ‘KLRG1’, ‘CD62L’, ‘MHCII’, and ‘MHCI’. B) Uniform manifold approximation projection plots show the distribution of cells based on relative surface protein expression. N=6 mice at 3-months old and N=5 mice at 18-months of age. n=3,776,804 total cells. All mice are female C57BL/6N.

We then plotted the young and aged individual marker expression as a dot heatmap by group (Figure 4A) and by individual animal (Figure 4B). We found that CD62L expression was significantly higher in young spleen T cells while the expression of CD44, CD49d, PD1, and CXCR3 were greater in aged splenic T cells (Figure 4B).

**Figure 4:**
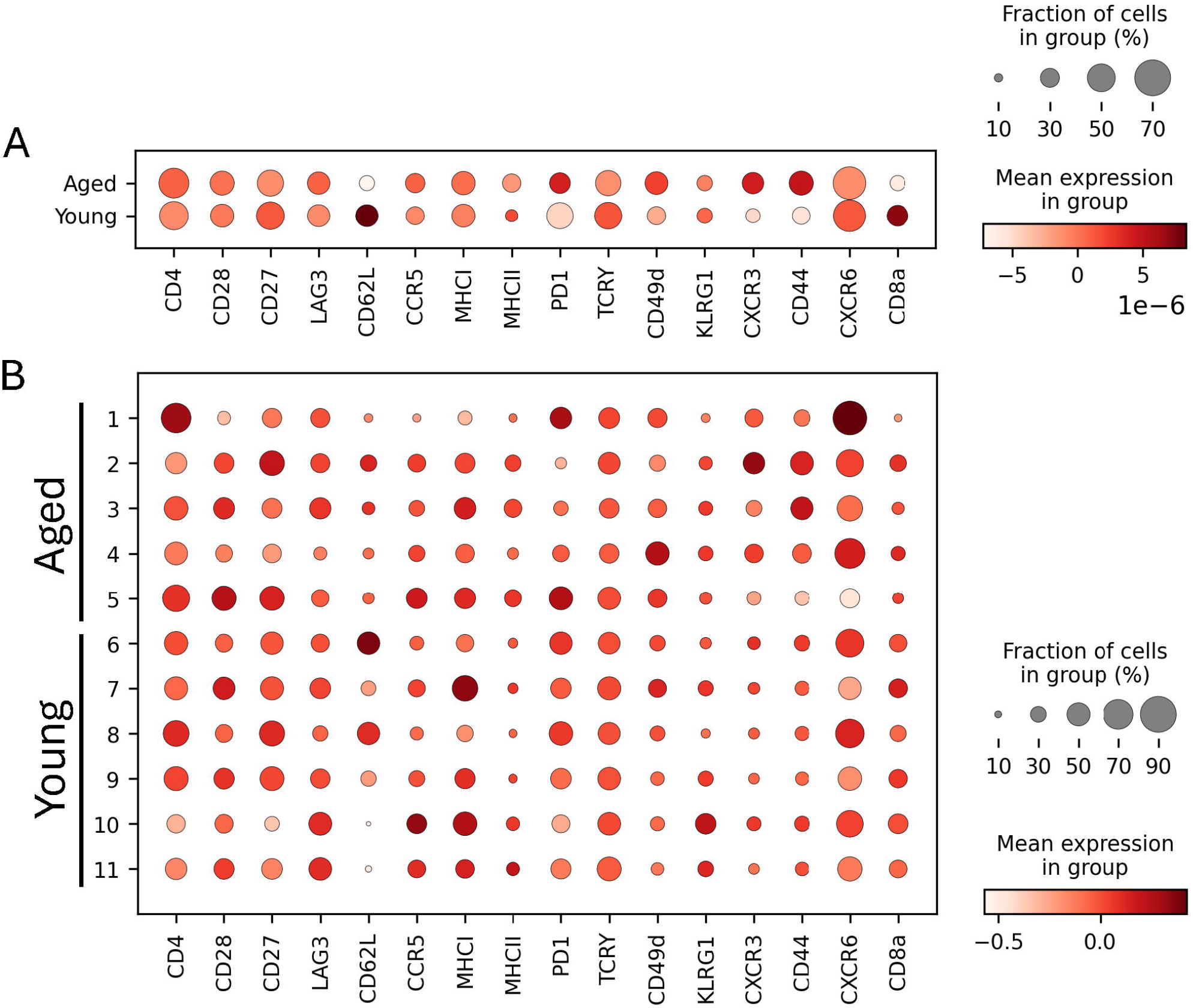
Expression of surface protein markers differs in young and aged mice. Dot-heatmap plots showing the fraction of cells expressing a marker and mean protein expression o 16 immune markers (CD49d, CXCR3, CCR5, CXCR6, CD44, CD8a, CD4, CD27, TCRY, CD28, PD1, LAG3, KLRG1, CD62L, MHCII, MHCI) by age group (A) and in individual animals (B). N=6 mice at 3-months old and N=5 mice at 18-months of age. n=3,776,804 total cells. All mice are female C57BL/6N.

We next quantified the total number of cells in each cluster by age group (Figure 5A) and frequency of each cluster as a percentage of all CD3e^+^ T cells (Figure 5B). Cluster 0 was the largest cluster and was split evenly between young and aged mice; however, clusters 1, 3, and 6 were increased in aged mice (Figures 5A and B). Clusters 2 and 7 were primarily present in young animals (Figures 5A and B). These data largely corroborate and provide quantification for the data shown in Figure 2. We also show quantification of these data by stacked barplot for total cell number (Figure 5C) and frequency (Figure 5D) by age group.

**Figure 5:**
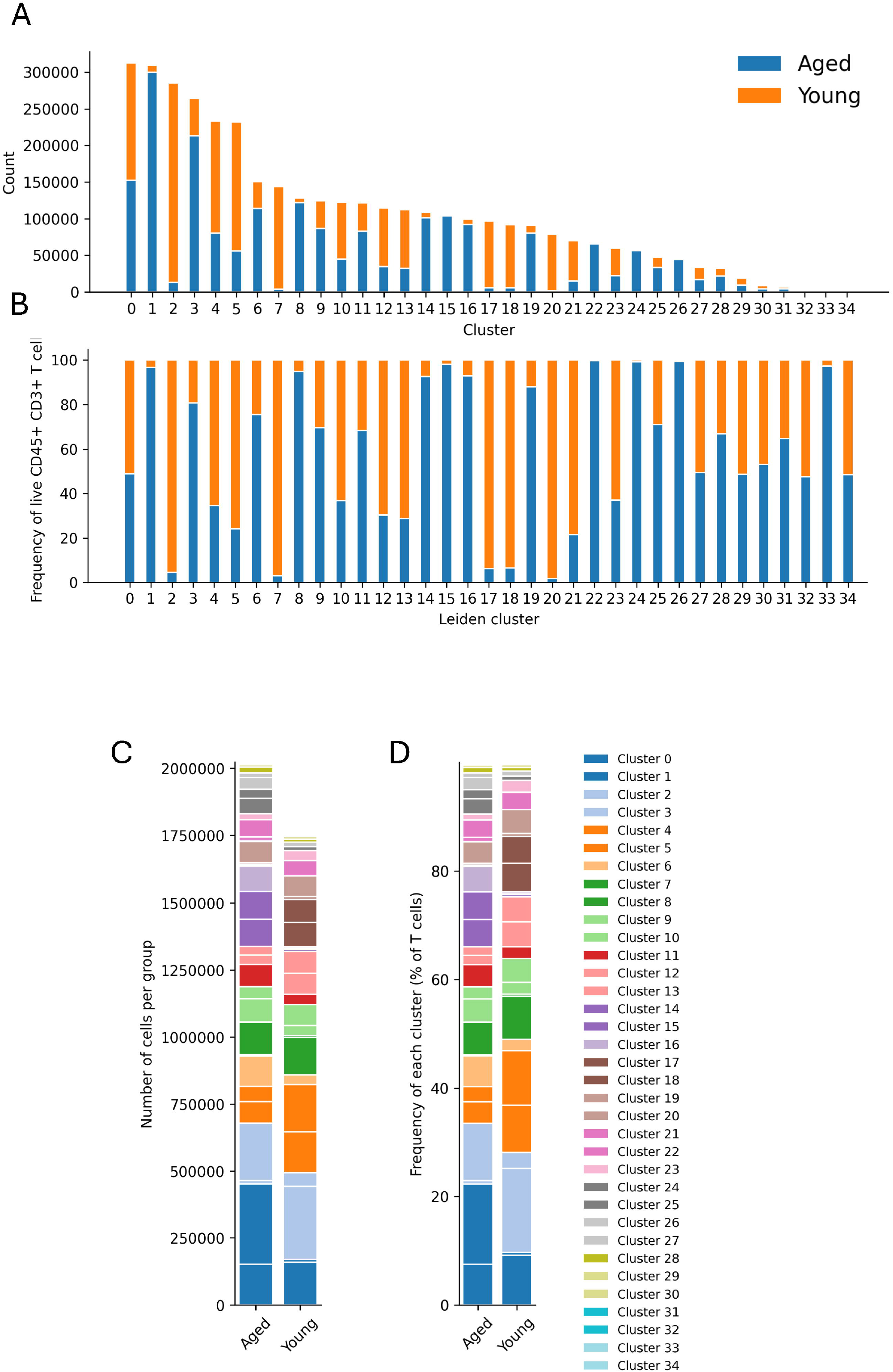
Quantitative distribution of T cells across leiden clusters by age group. Individual stacked bar plots for each cluster by age group show the total number (A) and frequency of CD3e^+^ T cells (B) of immune cells across 34 leiden clusters. These data are also shown as a combined stacked bar plot for both the total number of cells (C) and frequency of CD3e^+^ T cells (D). N=6 mice at 3-months old and N=5 mice at 18-months of age. n=3,776,804 total cells. All mice are female C57BL/6N.

Finally, our analysis pipeline includes export of total cell count, frequency, and median fluorescence intensity from individual samples which we imported into GraphPad Prism software for analysis and plotting. We found several statistically significant differences in the frequency of T cells among clusters between young and aged groups across CD8^+^ T cells, CD4^+^ T cells, and γδT cell clusters. Within the CD8^+^ T cell clusters, we found that clusters 4, 17, 18, and 20 were statistically significantly increased in young animals while clusters 14 and 19 were increased in aged mice (Figure 6A). We identified more clusters of CD4^+^ T cells compared to CD8^+^ clusters and more of these were significantly altered by age group. CD4^+^ T cell clusters 2, 5, 7, and 10 were greater in young animals while clusters 1, 3, 6, 8, and 16 were increased in aged animals (Figure 6B). We also identified 5 clusters of γδT cells and other unknown T cell subsets. Of these, clusters 12 and 23 were increased in young animals while cluster 15 was significantly increased in aged mice (Figure 6C).

**Figure 6:**
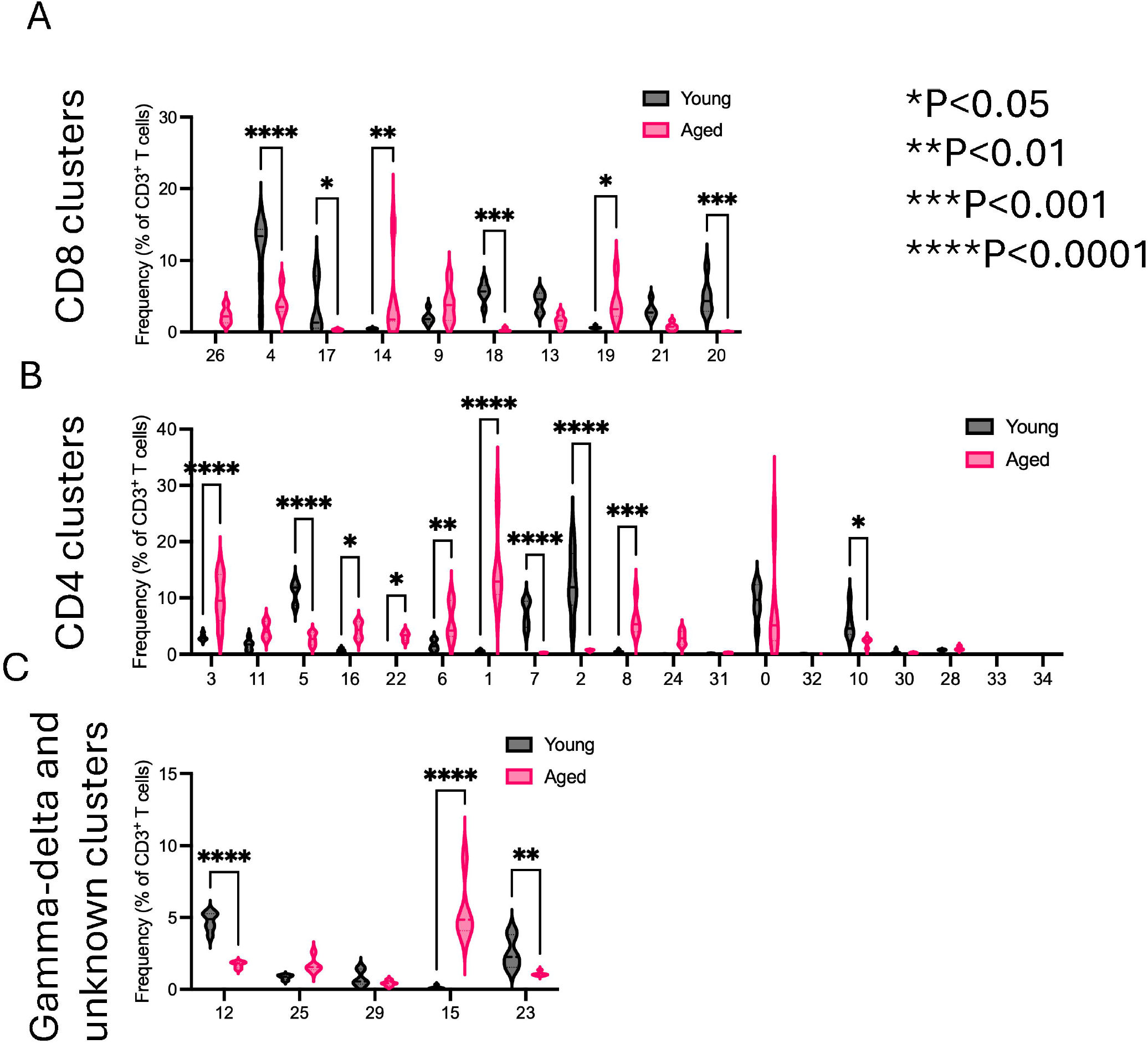
T cell clusters are statistically significantly different by age group. Frequency of each of the leiden T cell clusters were exported from our analysis pipeline and statistical analysis was performed in GraphPad Prism software using one-way ANOVA. Data are shown for CD8^+^ T cell clusters (A), CD4^+^ T cell clusters (B), and γδ T cell clusters (C). N=6 mice at 3-months old and N=5 mice at 18-months of age. n=3,776,804 total cells. All mice are female C57BL/6N. Data are mean ± quartiles. *P<0.05, **P<0.01, ***P<0.001, ****P<0.0001.

Using the marker protein expression from Figure 3A and which clusters were significantly different based on age in Figure 6, we sought to determine whether it was possible to manually gate on cells expressing markers in a bi-axial manner. Cluster 6 is a CD4 T cell cluster characterized by high CD27 and PD-1 expression and low expression of CD62L, CCR5, CXCR3, and KLRG1. We manually gated on CD27+ and CD62L-cells, then PD-1+ and CXCR3-cells, then CCR5- and KLRG1-cells to isolate the potential Cluster 6 population (Supplemental Figure 5A). We show a representative cell population by manual gating (Supplemental Figure 5B) and by our analysis pipeline (Supplemental Figure 5C) for Young Sample #4 and Aged Sample #1. We then quantified Cluster 6 from our analysis pipeline and manual gating and demonstrated that Cluster 6 was significantly increased in the aged samples compared to young. By leiden algorithm, Cluster 6 comprised ~5% of aged CD3^+^ T cells and ~10% by manual gating (Supplemental Figure 5D).

## Discussion

In this paper, we show an example of spectral flow cytometry using 22 colors to phenotype T cell subsets. Spectral flow cytometry is similar in some ways to other high dimensional single-cell analyses like cyTOF.^34–38^ Significant differences between the two are that spectral flow cytometry uses fluorochrome-labeled antibodies in contrast to heavy metal-labeled antibodies in cyTOF.^37^ Spectral flow cytometers can analyze cells at a similar rate as conventional flow cytometers (10 to 5,000 cells/second) which contrasts with cyTOF which analyzes cells at a slower rate (10-1,000 cells/second).^3,39–42^ Thus, a greater number of cells can be analyzed using spectral flow cytometry compared to cyTOF. For example, here we performed spectral flow cytometry on 3 million cells in contrast to published cyTOF datasets that contain between 1,000 and 500,000 cells.^30,42,43^ The availability of fluorochrome-labeled antibodies is a significant advantage of spectral flow cytometry compared to cyTOF in terms of cost and ease of implementation. A limitation of using spectral flow cytometry is that data analysis becomes increasingly more cumbersome with each additional antibody added to the panel compared to conventional flow cytometry where the number of antibodies is limited by spectral overlap.

Single-cell analyses including spectral flow cytometry, cyTOF, and single-cell RNA sequencing are high-dimensional data structures. Several algorithms and plotting functions have been developed to reduce dimensionality in order to show high dimensional data in 2 dimensions for visual representation. For flow cytometry data, popular clustering techniques include FlowSOM^29^, PhenoGraph^30^, and X-shift^31^. PhenoGraph is based on the Louvain algorithm^44^; however, this was shown to yield badly connected or disconnected communities. The Leiden algorithm was developed to improve upon the Louvain algorithm to identify better-connected communities.^27^ X-shift relies on k-nearest neighbors and FlowSOM relies on a self-organizing map (SOM) with hierarchical clustering. Here, we developed a spectral flow cytometry analysis pipeline based on existing cutting-edge single-cell analyses using the Leiden algorithm from the Scanpy package.^26^

Analyzing spectral flow cytometry data using conventional gating strategies with two antibodies per analysis step in a biaxial manner becomes difficult with large panels. For example, if attempting to phenotype CD8^+^ T cell subsets with a conventional gating strategy, one would typically gate on live cells, then CD45^+^ cells, CD3e^+^ T cells, then CD8a^+^ CD4^-^ cells.^18,45^ Once the CD8^+^ T cell population is isolated, five markers will have been used in conventional flow cytometry. In some cases, dump channels for other cell types like B cells (B220), myeloid cells CD11b, CD11c, and Ly-6g may be used. Typical gating strategies for murine CD8 T cells then employ markers to differentiate naïve, effector memory, and central memory cells. At minimum, this means that CD44 and CD62L will be used to differentiate these three populations.^17^ Sometimes additional markers like markers for T cell exhaustion (i.e. PD-1) and chemokine receptor markers like C-C Chemokine Receptor 5 (CCR5) will be used to determine more nuanced phenotypes of these three populations. Using these four additional markers after isolating the CD8^+^ cell population limits the decision of which markers to choose for the first two antibodies, which are typically CD44 and CD62L, followed by the phenotypic markers PD-1 and CCR5.^17^ However, with spectral flow cytometry, there may be an additional 14 or more markers after isolating the CD8^+^ population. In this situation, one may still decide to drill down into naïve, effector memory, and central memory cells in a biaxial manner first with CD44 and CD62L. After this, there are 12 phenotypic markers, which in our example here included CD27, CD28, PD1, CCR5, CXCR3, CXCR6, KLRG1, LAG3, CD49d, MHC-I, MHC-II, and TCR-γ. It is difficult to determine what order to examine these markers in if they must be examined sequentially in a biaxial manner in two dimensions. This is why we sought to examine these data using tools designed for high dimensional, large data matrices. Single-cell RNA sequencing has the most advanced data analysis tools in this research area. The major differences between scRNAseq and spectral flow cytometry are that scRNAseq data are typically in a large, sparse matrix of data where around 5,000-40,000 cells have 10,000-25,000 genes expressed. In scRNAseq, most genes are not expressed in any given cell, which results in a sparse data matrix. In contrast to this, spectral flow cytometry data may include 10,000-100,000,000 cells where each cell expresses between 15-50 proteins. This results in a dense data matrix but with a similar number of datapoints as scRNAseq. For this reason, we wanted to use single-cell RNA sequencing analysis tools to analyze spectral flow cytometry data.

To demonstrate the utility of our spectral flow cytometry antibody panel and our analysis pipeline, we examined young, 3-month, and old, 21-month, splenocytes from female C57BL/6N mice. We used principle components analysis with UMAP to reduce dimensionality for 2-dimensional plotting.^28^ We used the Leiden^27^ algorithm for clustering to identify 34 unique T cell clusters. Caution should be used when using extreme dimensional reduction techniques on entire datasets.^46^ Especially when inferring biological significance from the distances between clusters or cells in UMAP images. Our pipeline uses principal components analysis to reduce dimensionality to increase computational efficiency; however, this also results in some loss of high dimensional relationships. If computational efficiency is not limited or smaller datasets are used, the nearest neighbors and Leiden algorithm can be applied directly to the high-dimensional data in our pipeline with a single word change in the code.

In our analysis, relying on the leiden clusters, we identified four clusters of naïve CD8 T cells characterized as CD62L^+^ and CD44^-^. Cluster 17 was significantly enriched in young spleen whereas cluster 26 was almost exclusively in old splenocytes, although not statistically significant. Cluster 9 was split more evenly between both young and old splenocytes with higher expression of CD44 compared to Cluster 7, suggesting these may be activated or an early central memory population.^47^ Cluster 14 is more aligned with the classic central memory population and is significantly enriched in the aged samples.^48^ Cluster 19 expressed CD44 but not CD62L suggesting it is an effector memory population. Cluster 19 also expressed chemokine receptors CXCR3 and CXCR6, integrin CD49d, and checkpoint molecule, PD-1. This suggests this population may be exhausted and prone to tissue residence.^49–51^ Clusters 18 and 20 had low expression of CD62L and CD44, suggesting these may be effector subtypes or in a transitional state.

Our analysis also demonstrated that there were significant age-related differences in γδ T cell clusters.^52,53^ Both clusters 12 and 23 were statistically significantly increased in young splenocytes, but cluster 15 was significantly increased in 21-month old splenocytes. Clusters 12 and 23 were primarily differentiated by activation markers where cluster 12 expressed markers of naïve T cells like CD27 and CD62L and cluster 23 had more cells expressing activation markers like CD44, KLRG1, CCR5, and LAG3.^54–56^ Cluster 15, which was significantly enriched in old mice was very similar to cluster 23 except that it had little expression of LAG3 and almost no cells expressing CD44.

Regarding CD4^+^ T cells, Clusters 1, 3, and 8 were significantly increased in the old mouse splenocytes. Clusters 1 and 8 had high expression of CXCR6. They appear differentiated by CD27 expression in cluster 8 which was low in Cluster 1.^57,58^ A limitation of using only surface marker expression is that our data is descriptive. The antibody panel could be adapted for intracellular cytokine staining to inform on functional differences. A benefit of using surface protein expression is that we can develop panels for FACS to isolate cell populations and examine them in-depth using *in vitro* studies or adoptive transfer *in vivo* studies. Functional differences of the clusters we identified may be explored in follow-up studies.

## Acknowledgements

We thank the staff at the UAB Flow Cytometry and Single Cell Core laboratory.

## Footnotes

This work was supported by the National Institutes of Health, National Institute on Aging Grants R00AG068309 and Nathan Shock Center which is supported by the National Institute on Aging of the National Institutes of Health under award number P30 AG050886 (to D.J.T.). The UAB Flow cytometry core is supported by the Center for AIDS Research, AI027767 and The O’Neal Comprehensive Cancer Center, CA013148.

## Author Contributions

D.J.T. conceived and designed research. D.J.T., M.A.A., and D.V.III conceived of the figures and performed the analysis. D.J.T., M.A.A., and D.V.III drafted the manuscript. D.J.T., M.A.A., D.V.III, C.B., H.S., and H.T. edited, revised, and approved the final manuscript.

---

**Supplemental Figure 1:**
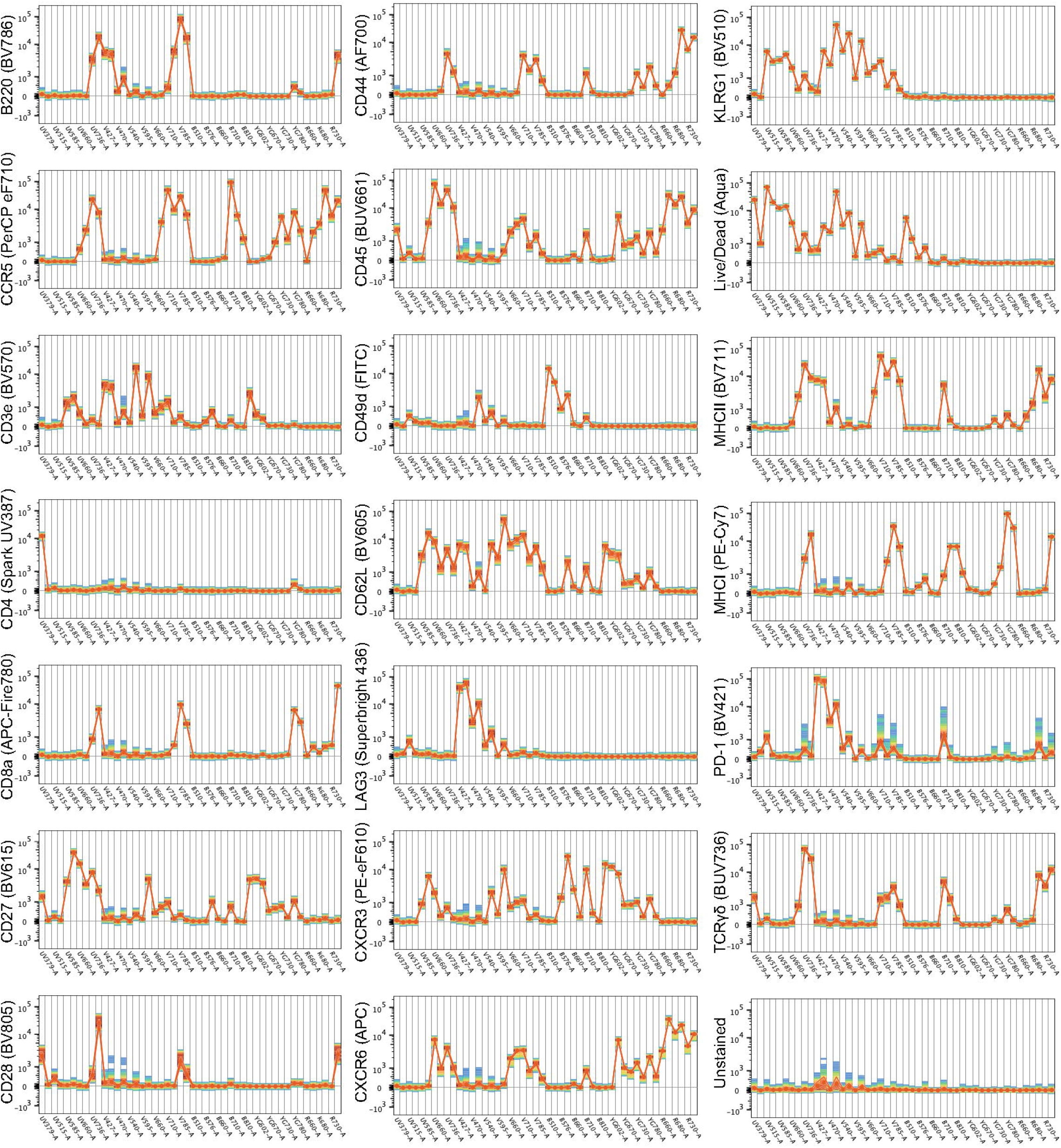
Spectral plots of single-antibody-stained bead samples including unstained negative control.

**Supplemental Figure 2:**
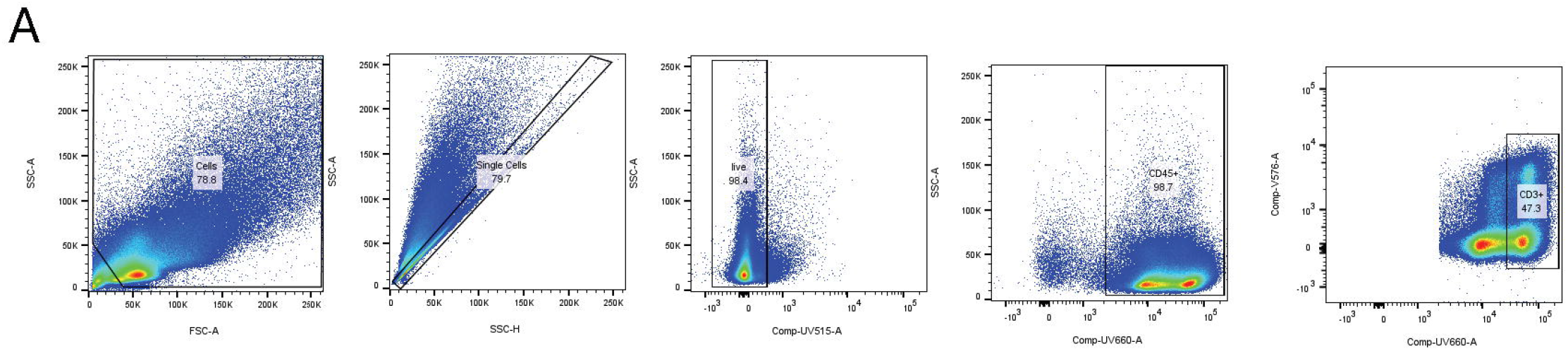
Conventional flow cytometry gating strategy to drill down onto CD3e^+^ cells prior to exporting scaled “.fcs” files for spectral flow cytometry.

**Supplemental Figure 3:**
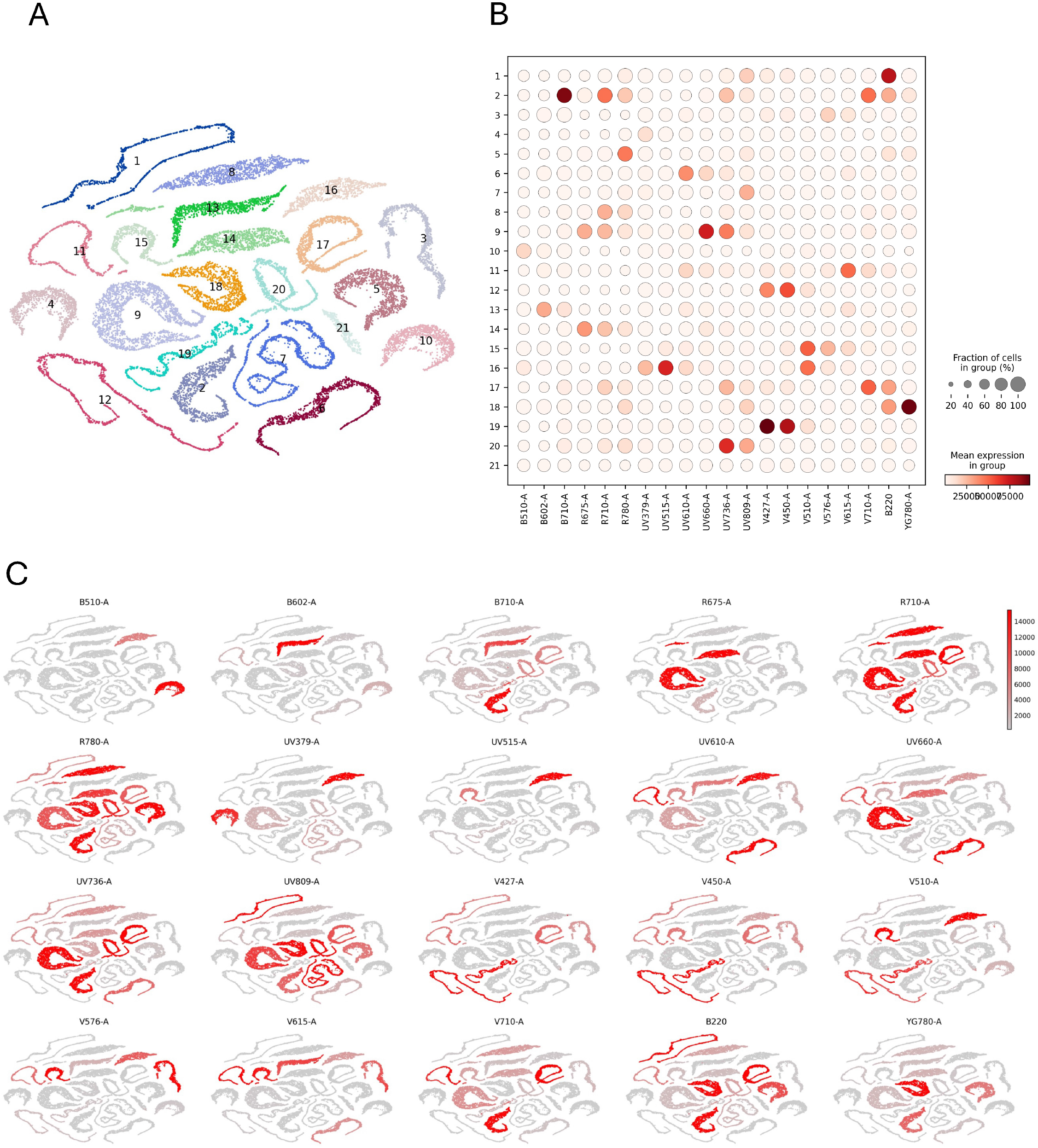
A) Uniform manifold approximation projection plot showing leiden clustering of single-antibody-stained bead samples and unstained sample with cluster number on the sample. B) Dot-heatmap of each of the 21 single-antibody-stained samples and unstained sample showing protein expression and the percent of expressing cells. C) Uniform manifold approximation projection plots show the distribution of single-antibody-stained beads by scaled surface protein expression.

**Supplemental Figure 4:**
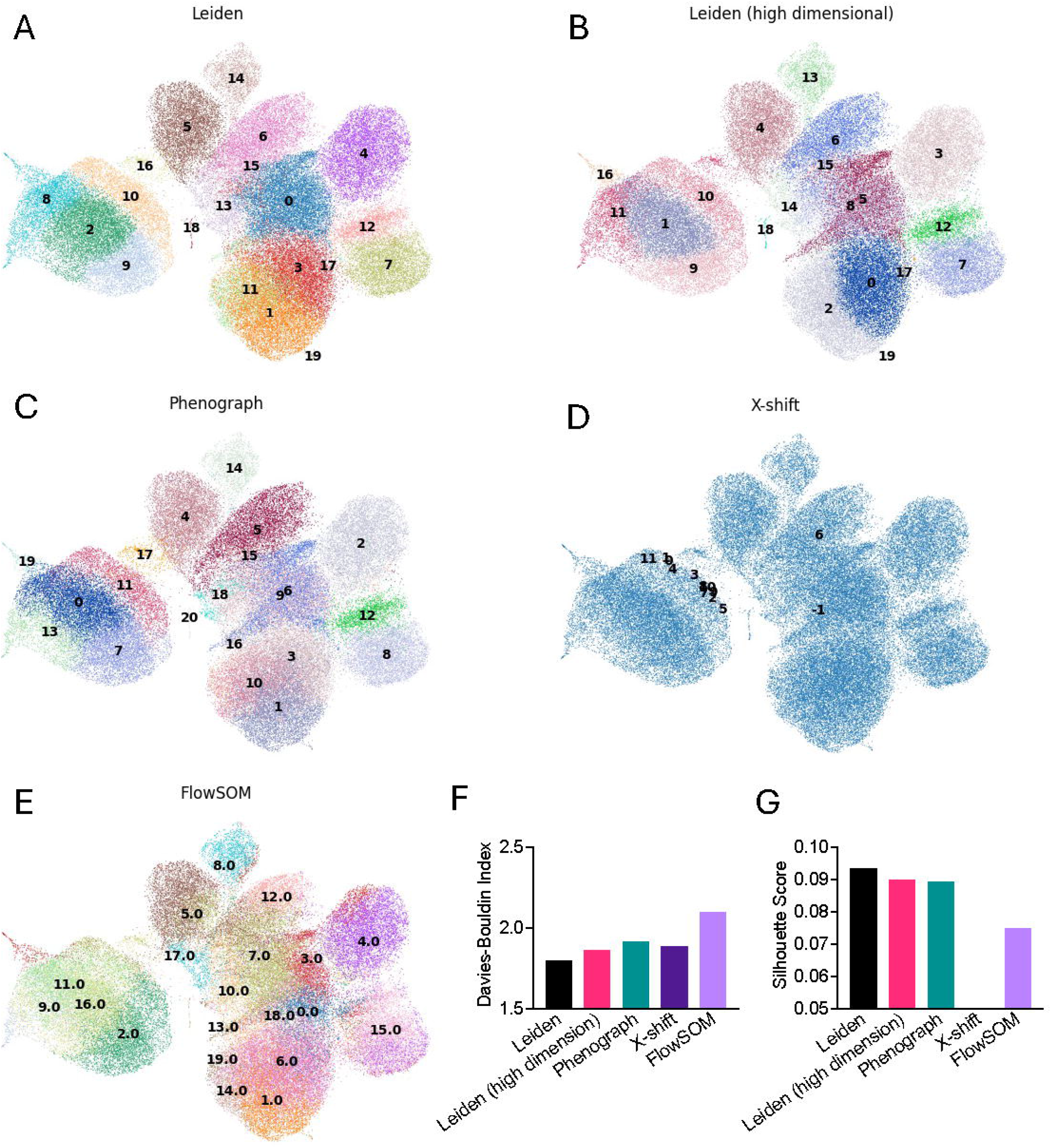
Brain T and B cells are processed, stained, and analyzed through our analysis pipeline by Leiden clustering (A), Leiden clustering on high dimensional data that did not undergo dimensional reduction by principle components analysis (B), Phenograph (C), X-shift (D), and FlowSOM (E). Davies-Bouldin Index (F) and Silhouette Score (G) were computed for each clustering method and compared. N=5 mice 3-months old that underwent sham surgery, N=5 mice 3-months old that underwent bilaterial carotid artery stenosis surgery, N=5 mice 18-months old that underwent sham surgery, and N=4 mice 18-months old that underwent bilaterial carotid artery stenosis surgery. n= 69,555 total cells. All mice are female C57BL/6N.

**Supplemental Figure 5:**
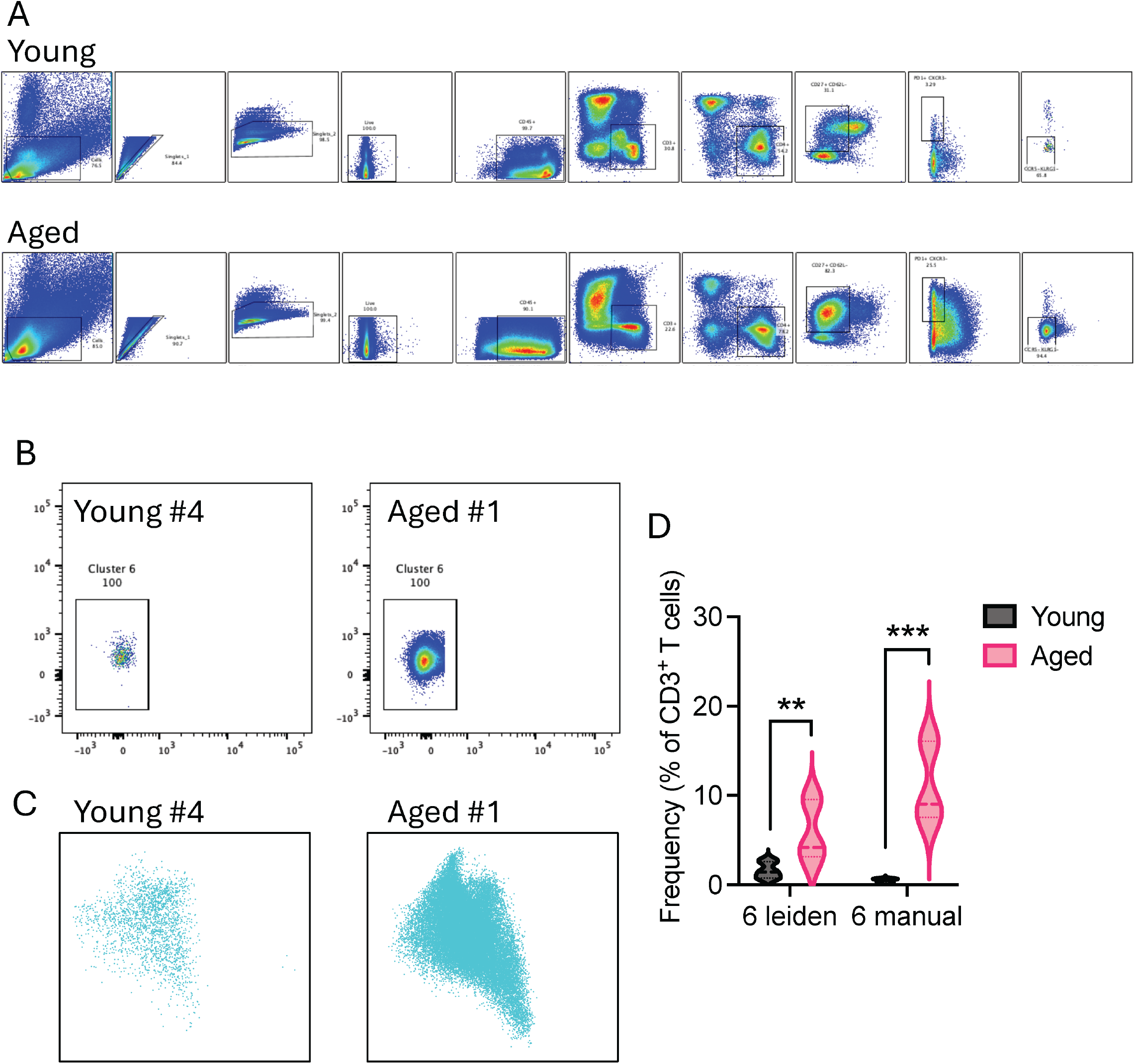
A) Gating strategy for Young sample #4 and Aged sample #1 to manually isolate Cluster 6 from the leiden clusters. B) Representative cell population of Cluster 6 for Young #4 and Aged #1 samples. C) Representative Cluster 6 by our analysis pipeline using Leiden algorithm for Cluster 6. D) Quantification of Cluster 6 identified by both leiden algorithm and manual gating. one-way ANOVA. N=6 mice at 3-months old and N=5 mice at 18-months of age. n=3,776,804 total cells. All mice are female C57BL/6N. Data are mean ± quartiles. *P<0.05, **P<0.01, ***P<0.001, ****P<0.0001.

